# Engineered bacteria that self-assemble “bioglass” polysilicate coatings display enhanced light focusing

**DOI:** 10.1101/2024.06.03.597164

**Authors:** Lynn M. Sidor, Michelle M. Beaulieu, Ilia Rasskazov, B. Cansu Acarturk, Jie Ren, Lycka Kamoen, María Vázquez Vitali, P. Scott Carney, Greg R. Schmidt, Wil V. Srubar, Elio A. Abbondanzieri, Anne S. Meyer

## Abstract

Photonic devices are cutting-edge optical materials that produce narrow, intense beams of light, but their synthesis typically requires toxic, complex methodology. Here we employ a synthetic biology approach to produce environmentally-friendly, living microlenses with tunable structural properties. We engineered *Escherichia coli* bacteria to display the silica biomineralization enzyme silicatein from aquatic sea sponges. Our silicatein-expressing bacteria can self-assemble a shell of polysilicate “bioglass” around themselves. Remarkably, the polysilicate-encapsulated bacteria can focus light into intense nanojets that are nearly an order of magnitude brighter than unmodified bacteria. Polysilicate-encapsulated bacteria are metabolically active for up to four months, potentially allowing them to sense and respond to stimuli over time. Our data demonstrate that engineered bacterial particles have the potential to revolutionize the development of multiple optical and photonic technologies.

**Significance Statement:** In this work, we apply the principles of synthetic biology to create living optical devices. Utilizing the ability of sea sponges to polymerize bioglass from silica precursors in the ocean water using only a single enzyme, silicatein, we have fused this same enzyme to the surface of *Escherichia coli* bacterial cells. The modified bacteria can polymerize a layer of bioglass at their surface. This bioglass shell allows the bacteria to act as engineered optical devices that are able to scatter high intensity, focused light while also surviving for several months, opening the door to a wide range of sense-and-respond applications.

**Classification:** Biological Sciences, Applied Biological Sciences

## Introduction

In nature, organisms have evolved innate abilities to produce multifunctional structures with complex compositions and advanced optical properties. These biologically-produced structures have vast potential to be used to design and produce new optical materials and devices using ecologically friendly manufacturing methods. In particular, certain biological structures show great promise as next-generation microlenses and optical microparticles. Microscopic organisms, cells, and materials produced by living organisms have already been shown to exhibit unique and beneficial properties for manipulating light. These “bio-microlenses” have been demonstrated to act as super-resolution magnifiers (*1*), to enhance upconversion in fluorescence (*2*), to sense light direction (*3*), to enable sub-diffractive focusing (*4*), and to act as waveguides (*5*). In addition, cell-based bio-microlenses are similar in size and shape to photonic structures that are used for subwavelength microscopy and the formation of photonic nanojets (*6–11*). However, in order to tailor biological particles to specific applications, we need the ability to tune their optical properties.

Synthetic biology offers a path to engineer single cells by altering their size, length, shape, and refractive index, all of which can be used to optimize their ability to act as bio-photonic microlenses. Synthetic biology approaches can also be used to combine unique optical functions of different organisms. Many aquatic organisms, including sea stars and sponges, possess the ability to synthesize natural structures that perform both optical and structural functions. The arms of the brittlestar are coated in light-sensitive calcite plates, which provide protection and also act as highly efficient light-capturing microlens arrays (*12, 13*). Similarly, hexactinellid sponges create silica spicules that are responsible for their structural stability and also display waveguide properties (*14, 15*). Siliceous sponges deposit silica into needle-like spicules, which is accomplished by polymerizing silica into polysilicate, also known as “bioglass,” using a unique silicatein enzyme (*16–20*). Silicatein-catalyzed silica deposition requires only a single gene to be expressed and can be performed at physiological temperature, pressure, and pH without the use of harsh or toxic chemicals (*21, 22*). Silicatein is thought to be the only natural biomineralizing enzyme (*23*), and its ability to fabricate polysilicate structures offers a powerful addition to the synthetic biology toolbox.

In this article, we use multiphysics modeling to demonstrate that polysilicate-encapsulated bacterial cells are predicted to have enhanced abilities to scatter and focus light into photonic nanojets. We then demonstrate for the first time that engineered *Escherichia coli* bacteria that express surface-displayed silicatein enzymes from sea sponges *Tethya aurantia* and *Suberites domuncula* can coat themselves in a layer of polysilicate. We observe that these cells scatter and focus light in a manner that resembles the modeled data. Using a fluorescent probe, we show that the polysilicate-encapsulated bacterial cells, when exposed to planar illumination, create intense beams of focused light that form a peak of intensity outside the cell. In contrast, wild-type bacteria create much weaker beams that peak at the cell surface. Remarkably, polysilicate- encapsulated cells survive for up to four months post-encapsulation and are still able to scatter light with a comparable focal peak after they become metabolically inactive. These optically- tuned bacteria are the first example of biologically engineered microlenses, and they have the potential to create new biologically-active devices with controllable properties for optimizing optical performance across a variety of applications including advanced biosensing, super-resolution imaging, Raman scattering, nanolithography, and photovoltaics (*3, 4*).

## Results

### Prediction of enhanced nanojets

We first set out to test whether a thin polysilicate shell would be expected to have a measurable impact on the ability of a bacterial cell to focus light. We therefore simulated the passage of light through a typically-sized *Escherichia coli* bacterial cell surrounded by a uniform coating of polysilicate using the finite-difference time domain method (Supp. Figure 1A-B), a method that has previously been used to model photonic nanojets and microspheres (*24, 25*). The electric field intensity throughout the simulation region is a superposition of incident and scattered light (*26*), allowing us to create intensity maps showing the brightness of the light in the vicinity of the bacterial cells (Figure 1A-B). At an angle of incidence of 0°, the intensity maps demonstrated that the encapsulated cells produced a bright central beam of scattered light, as well as less-intense peripheral beams (Figure 1A). Wild-type cells were modeled identically to the encapsulated cells but without a polysilicate coating, and the intensity maps of the wild-type cells showed a beam that was less intense compared to the encapsulated cells (Figure 1B).

**Fig.1:**
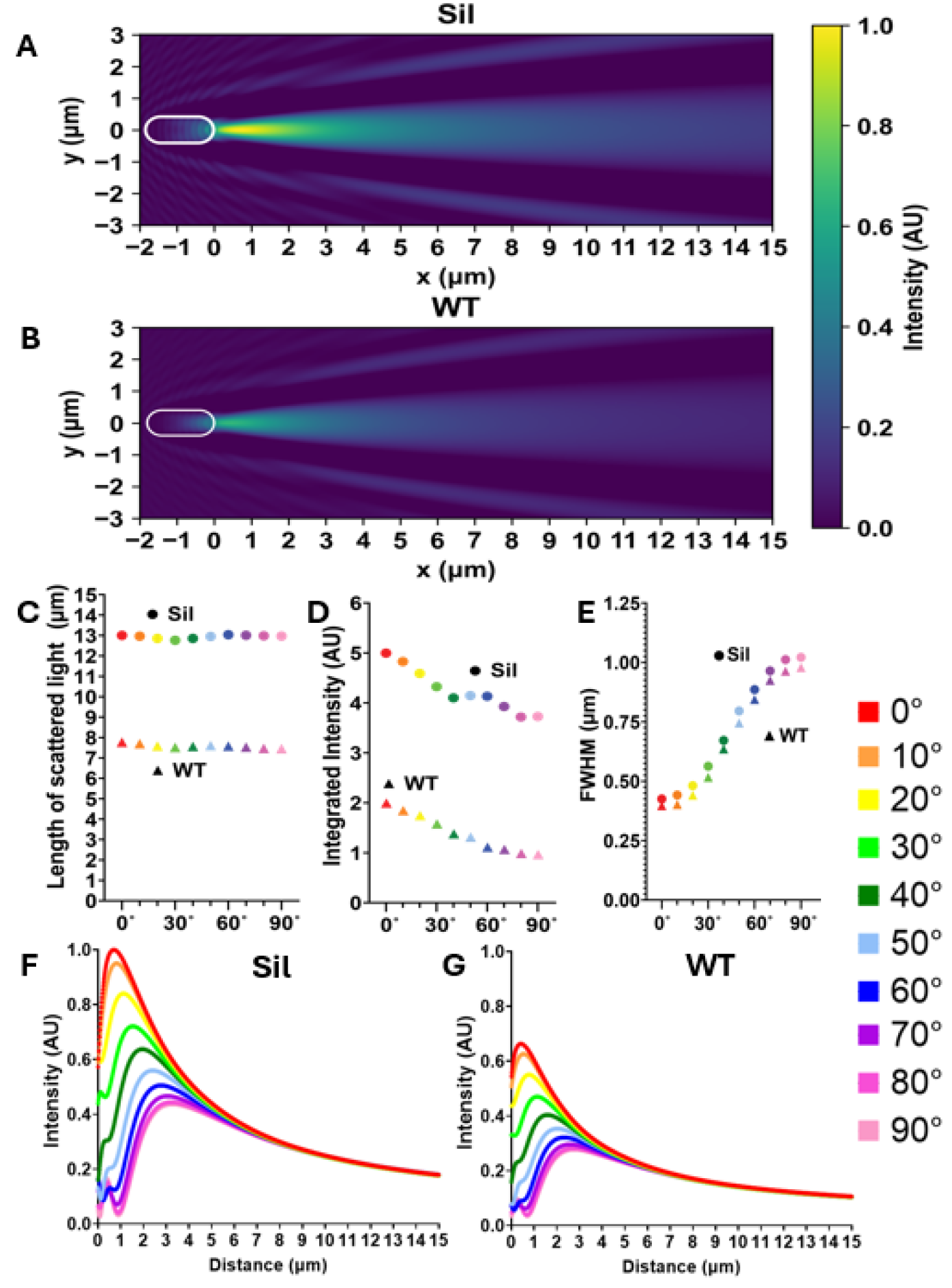
Mathematical simulations predict that polysilicate-encapsulated bacteria can produce photonic nanojets. (A-B) Intensity map showing the light scattered by a modeled polysilicate-encapsulated bacteria cell (Sil) (A) and a modeled wild-type cell (WT) (B). (C-G) Each color corresponds to a different angle of the cell in relation to the incident light, where (C) is the length of scattered light, (D) is the normalized integrated intensity of the scattered light, and (E) is the FWHM. (F-G) Normalized intensity of the scattered light as a function of distance from the edge of the cell for different angles of incident light for polysilicate-encapsulated cells (Sil) (F) and wild-type cells (WT) (G).

Simulations were next performed for bacterial cells at a range of angles relative to the incident light, to capture the effect of cell orientation on light scattering. For both wild-type and polysilicate-coated simulated cells, the intensity was calculated for incident light angles between 0°-90° at 5° or 10° increments. In order to quantify the length and intensity of the central scattered beam, we determined the boundary where the beam intensity exceeded the background intensity by 30%. This calculation allowed us to define the length and integrated intensity of the scattered beam (Figure 1C-D). Although the polysilicate layer only modestly increased the diameter and average index of refraction of the cells, the scattered beams of light were significantly longer and brighter compared to the modeled wild-type cells. To determine the width of the beam, we calculated the full width at half maximum (FWHM) of the central peak after subtracting the background intensity (Figure 1E, Supp. Figure 1C-D). The narrowest beams had widths below 0.4 µm (0.8 λ, where λ is the wavelength of illuminating light). The encapsulated cells produced beams that were focused deeper into solution and were moderately broader (Supp. Figure 1E-F). The focal shift due to encapsulation ranged from 0.28 µm deeper at 0° incidence up to 0.49 µm deeper at 90° incidence. The angle of the modeled cells relative to the incoming light was seen to have an effect on both the intensity of the scattered light and the location of the focal peak relative to the bacterial cell, with larger angles resulting in lower-intensity scattered light with a peak farther away from the cell body (Figure 1F-G). Taken together, these modeling data indicate that polysilicate-coated bacterial cells are expected to generate significantly brighter and longer photonic nanojets compared to wild-type bacterial cells.

### Construction of silicatein-expressing strains

In order to construct bacteria strains that express silicatein enzymes on the cell surface, where they can interact with silica precursor molecules in solution and biomineralize a layer of polysilicate bioglass, silicatein enzymes from *Tethya aurantia* or *Suberites domuncula* sea sponges were fused to a bacterial outer membrane protein (outer membrane protein A [OmpA] (*27*) or ice nucleation protein (*28*) [INP]). Constructs containing INP-*T. aurantia* silicatein were not able to be created either though gene synthesis or traditional restriction enzyme-based assembly. Expression of an INP-*S. domuncula* silicatein construct in *E. coli* resulted in high rates of abnormal cell elongation morphology following the polysilicate encapsulation protocol and showed a varying pattern of polysilicate staining with Rhodamine123. Therefore, INP-silicatein fusions were not used further. To achieve multiple types of induction control, OmpA-silicatein constructs were cloned into several inducible plasmids (pBAD33, pTrc99a, pRHA109, pRHA113, and pBbS5a). Any constructs cloned into pTrc99a were lethal upon induction, and cloning into pRHA109 was repeatedly unsuccessful. OmpA-silicatein constructs cloned into pBAD33 showed two distinct cell populations after induction: cells that demonstrated successful peripheral Rhodamine123 staining and cells that showed low-intensity, homogeneous staining, likely indicative of non-encapsulated cells; these strains were not used further. The *E. coli* strain expressing pBbS5a OmpA-*Tethya aurantia* silicatein (TaSil) showed consistent peripheral Rhodamine123 staining patterns and normal cell morphology and is used throughout this work. The only pRHA113 construct that was able to be synthesized was OmpA-*S. domuncula* silicatein (SdSil), which is used in this work as well. We attribute the variable success of these constructs to the overexpression of the outer membrane proteins, which can negatively impact cell viability, and the hydrophobicity of the silicatein enzyme.

### Silicatein localizes to cell borders

For our constructed strains that expressed OmpA-silicatein fusion proteins, we tested whether the silicatein enzyme was displayed on the surface of the *E. coli* cells after silicatein induction and incubation with orthosilicate, via performing immunofluorescence using an anti-silicatein antibody. Confocal imaging revealed that the immunofluorescence signal was localized to the bacterial cell surface on the cells expressing the OmpA-silicatein constructs (Figure 2A-B). The SdSil-expressing strain showed the localization of the enzyme to broad regions of the cell surface while the TaSil-expressing strain showed more punctate localization. The wild-type cells showed background levels of binding of the anti-silicatein antibody with no specific localization pattern (Figure 2C). Quantification of the immunofluorescence intensities revealed that the silicatein- expressing strains both showed significantly higher immunofluorescence than the wild-type control cells (Figure 2D). The differences in the immunofluorescence intensities and localization patterns between the TaSil- and the SdSil-expressing cells may be attributable to the fact that the anti-silicatein antibody was raised against spicules from *S. domuncula* sea sponges, which express a version of the silicatein enzyme that is approximately one-third smaller than the *T. aurantia* silicatein. Immunofluorescence on both the TaSil and SdSil strains after silicatein induction but not incubation with orthosilicate showed comparable antibody binding intensities and spatial distributions to those seen for the same strains incubated with orthosilicate, while the empty vector control strain showed little-to-no binding, similar to the wild-type strain (Supp. Figure 2A-D). These immunofluorescence results indicate that the engineered strains are able to express and display silicatein enzymes localized to their outer surfaces.

**Fig. 2:**
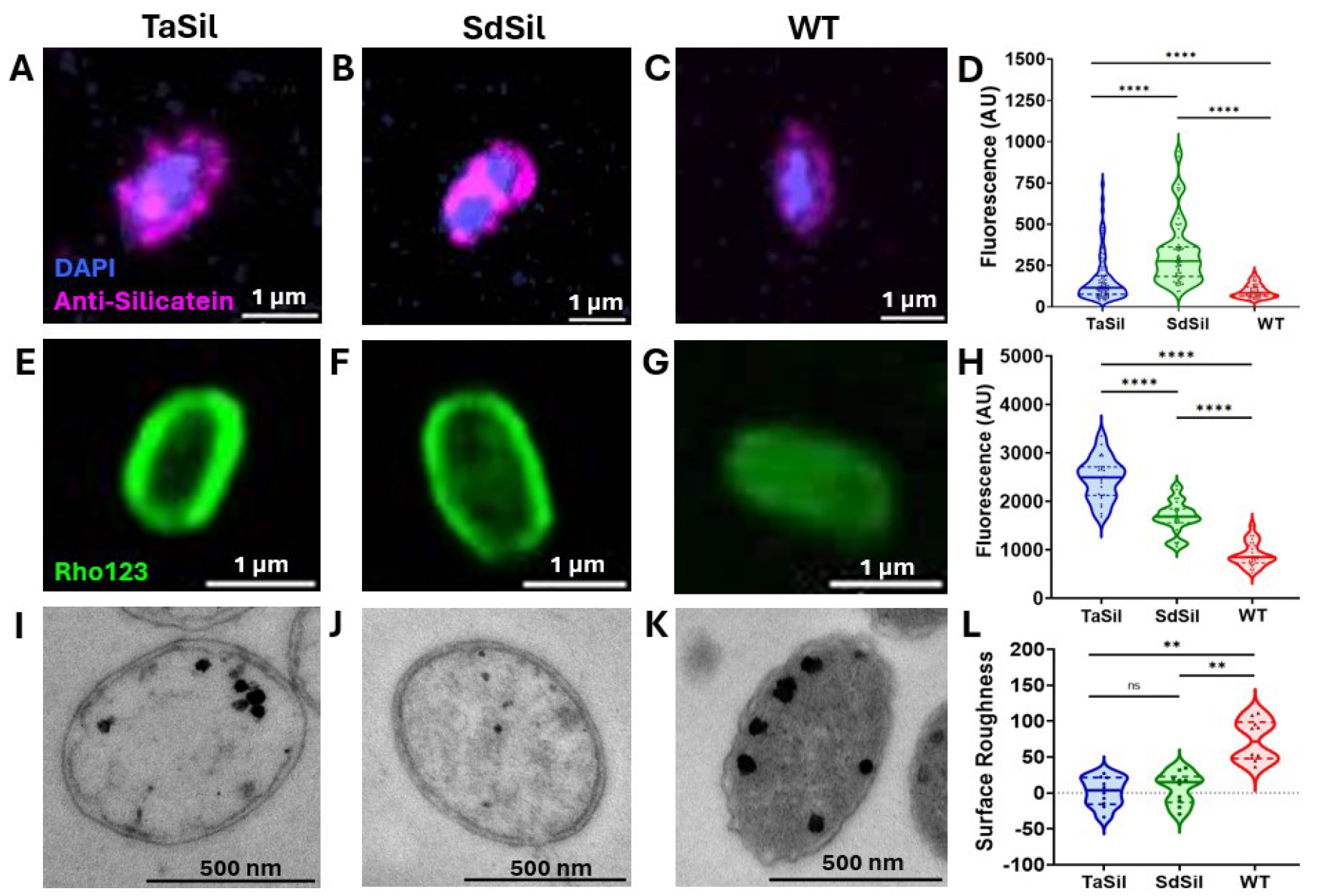
Silicatein-expressing cells display a smooth polysilicate border. (A-C) Immunofluorescence of silicatein-expressing (TaSil and SdSil) (A and B) and wild-type (WT) (C) strains following silicatein induction and incubation with orthosilicate. (D) Quantification of antibody fluorescence intensity of A-C (nTaSil=90, nSdSil=60, nWT=60). (E-G) Rhodamine123 staining of silicatein-expressing (E and F) and wild-type (G) strains. (H) Quantification of maximum Rhodamine123 fluorescence signal of E-G (n=20). (I-K) TEM thin section imaging of silicatein-expressing (I and J) and wild-type (K) strains. (L) Quantification of surface roughness of I-K (n=10). **** P≤0.0001, ** P≤0.01, ns: not significant.

### Encapsulation with polysilicate

To determine whether the silicatein-expressing strains were able to mineralize a layer of polysilicate surrounding the cells, the strains were induced, incubated with orthosilicate, and stained with Rhodamine123 dye, which adsorbs to silicate materials (*7, 10*). Both of the silicatein-expressing strains showed a bright Rhodamine123 signal localized to the outer border of the cells, while wild-type cells showed dim, diffuse staining (Figure 2E-G, Supp. Figure 3A- C). The maximum intensity of the Rhodamine123 signal for individual cells was significantly higher for TaSil and SdSil strains than for wild-type cells (Figure 2H). Quantification of the ratio of the border-to-internal fluorescence signal for Rhodamine123-stained cells indicated that the Rhodamine123 signal was several times higher on the border of the TaSil and SdSil cells than in the interior of the cells (Supp. Figure 3D). The border-to-internal ratios of Rhodamine123 fluorescence for the TaSil and SdSil cells were significantly higher than the ratios for the wild-type cells, which showed a ratio close to 1. TaSil and SdSil strains that were induced but not incubated with orthosilicate, as well as the empty vector control strain, all showed dimmer staining phenotypes (Supp. Figure 4A-F) and significantly lower border-to-internal fluorescence ratios (Supp. Figure 4G) than the engineered strains that were incubated with orthosilicate. These results indicate that the expression of silicatein enzymes on the outer surfaces of the TaSil and SdSil *E. coli* cells allowed them to encapsulate themselves in a layer of polysilicate coating the outside of the cells.

To determine the longevity of the biomineralized polysilicate coating on the engineered cells, Rhodamine123 staining was repeated on strains that had been stored in buffer for extended periods of time following encapsulation with polysilicate. One-month-old and five-month-old encapsulated TaSil and SdSil cells continued to display bright staining on their outer surfaces and had a border-to-internal fluorescence ratio that was significantly higher than for the one-month-old and five-month-old wild-type cells (Supp. Figure 3D, 5A-C). The five-month cells showed higher border-to-internal fluorescence ratios than cells measured at earlier timepoints (Supp. Figure 3D, 5D-F), potentially due to increased Rhodamine123 staining in dead or metabolically inactive cells (*29*). These data indicate that the polysilicate layer surrounding on the engineered cells can be stable for several months following initial biomineralization.

### Encapsulated cells have smoother borders

To determine the cell morphology of the polysilicate-coated cells, transmission electron microscopy (TEM) was performed on thin sections of cells for each strain following silicatein induction and incubation with orthosilicate (Figure 2I-K). The polysilicate-encapsulated cells showed a smooth, non-ruffled cell border phenotype, while wild-type cells had a typical ruffled outer membrane phenotype (*30*). The cell perimeter values of the thin-sectioned cells were determined from the TEM images and used to calculate cell surface roughness values. The polysilicate-coated cells showed significantly smoother cell perimeters than the wild-type cells and empty-vector control cells (Figure 2L, Supp. Figure 6A-E). The surface roughnesses of the polysilicate-coated TaSil and SdSil cells were not significantly different from each other (Figure 2L, Supp. Figure 6E). Additionally, we observed that the wild-type cells displayed a darker, more electron-dense cell interior than the polysilicate-encapsulated strains. Quantification of the intensities of the cell interiors revealed that the polysilicate-coated strains were significantly less electron-dense than the wild-type cells (Supp. Figure 6F). These analyses indicate that polysilicate coating of the TaSil and SdSil cells results in the formation of a smooth outer cell surface.

### Detection of silica deposition

To analyze the silica composition on the surface of our silicatein-expressing cells, the strains were induced, incubated with orthosilicate, and analyzed via scanning electron microscopy-energy dispersive X-ray spectroscopy (SEM-EDS). SEM images of the morphology of the cells showed that the silicatein-expressing cells appeared to be fused together in a viscous layer at the high densities used for this experiment, consistent with previous reports (*7*) (Supp. Figure 7).

EDS spectra indicated the presence of silica in the TaSil and SdSil cells, which displayed significantly more silica than the wild-type cells (Figure 3). Incubation of the wild-type cells with orthosilicate did not result in the detection of a significantly different amount of silica than for wild-type cells without the added orthosilicate, indicating that the levels of silica measured for both wild-type samples represent the background level of detection for this assay. X-ray diffraction (XRD) analysis was also consistent with the presence of silica-containing minerals in our silicatein-expressing, polysilicate-encapsulated strains (Supp. Figure 8).

**Fig. 3:**
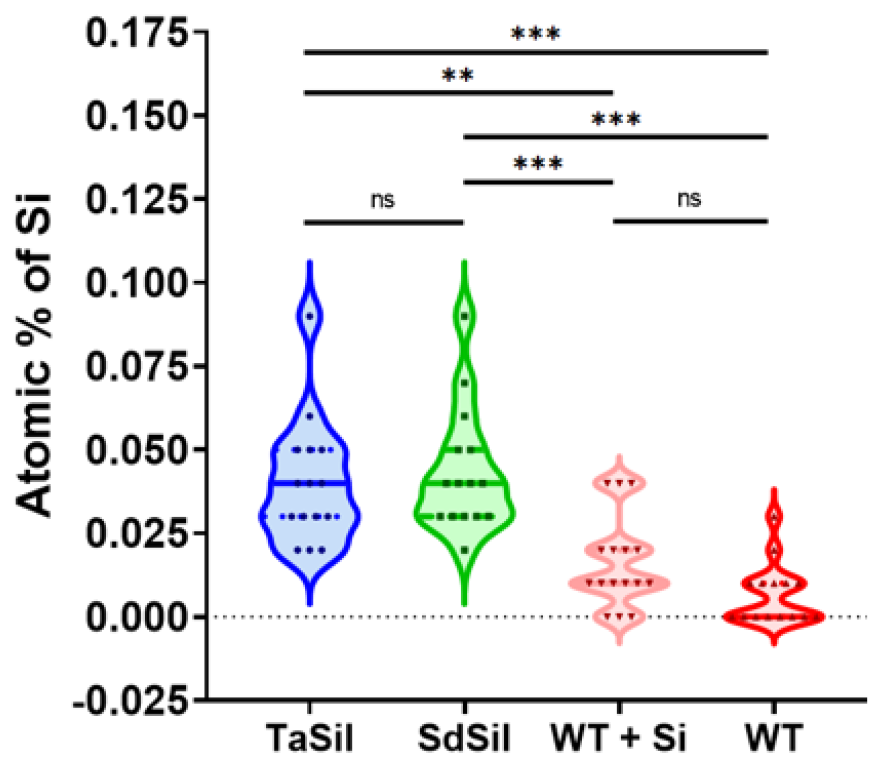
Silica detection on the surface of silicatein-expressing cells via SEM-EDS. Quantification of the atomic percent of elemental silica from EDS spectra readings of silicatein- expressing strains (TaSil and SdSil) following silicatein induction and incubation with orthosilicate, and the wild-type strain incubated (WT + Si) or not incubated (WT) with orthosilicate (nTaSil=15, nSdSil=15, nEV=13, nWT=15, nWT+silica=16). ** P≤0.01, ***P≤0.001, ns: not significant.

### Polysilicate-coated cells can generate photonic nanojets

To determine the ability of our polysilicate-encapsulated cells to scatter and focus light, we used a custom-built microscope (Supp. Figure 9) capable of illuminating the bacteria at incident angles ranging from -90° to 90°, a technique we term Multiple Angle Illumination Microscopy (MAIM). By adjusting the position of the illuminating beam at the back focal plane of a 1.49 NA oil-immersion objective (Figure 4A), we could continuously vary the illumination from vertical (i.e. epi-illumination) to horizontal before the beam is cut off by total internal reflection. Cells were imaged on an agarose pad, and the liquid and agarose around the cells contained the fluorescent dye Alexa-488, allowing us to directly visualize the size and shape of the scattered light above the coverslip. We continually recorded the fluorescence around the bacteria as the illumination angle was varied from -90° to +90°. The polysilicate-encapsulated cells were seen to scatter a bright jet of light that was most visible at near-horizontal illumination and showed close agreement with the scattering patterns that were predicted to be imaged by multiscale modeling (Supp. Figure 10). To ensure that each cell was exposed to the same set of illumination conditions, we recorded the scattered light across the full range of illumination angles (Supp.

**Fig. 4:**
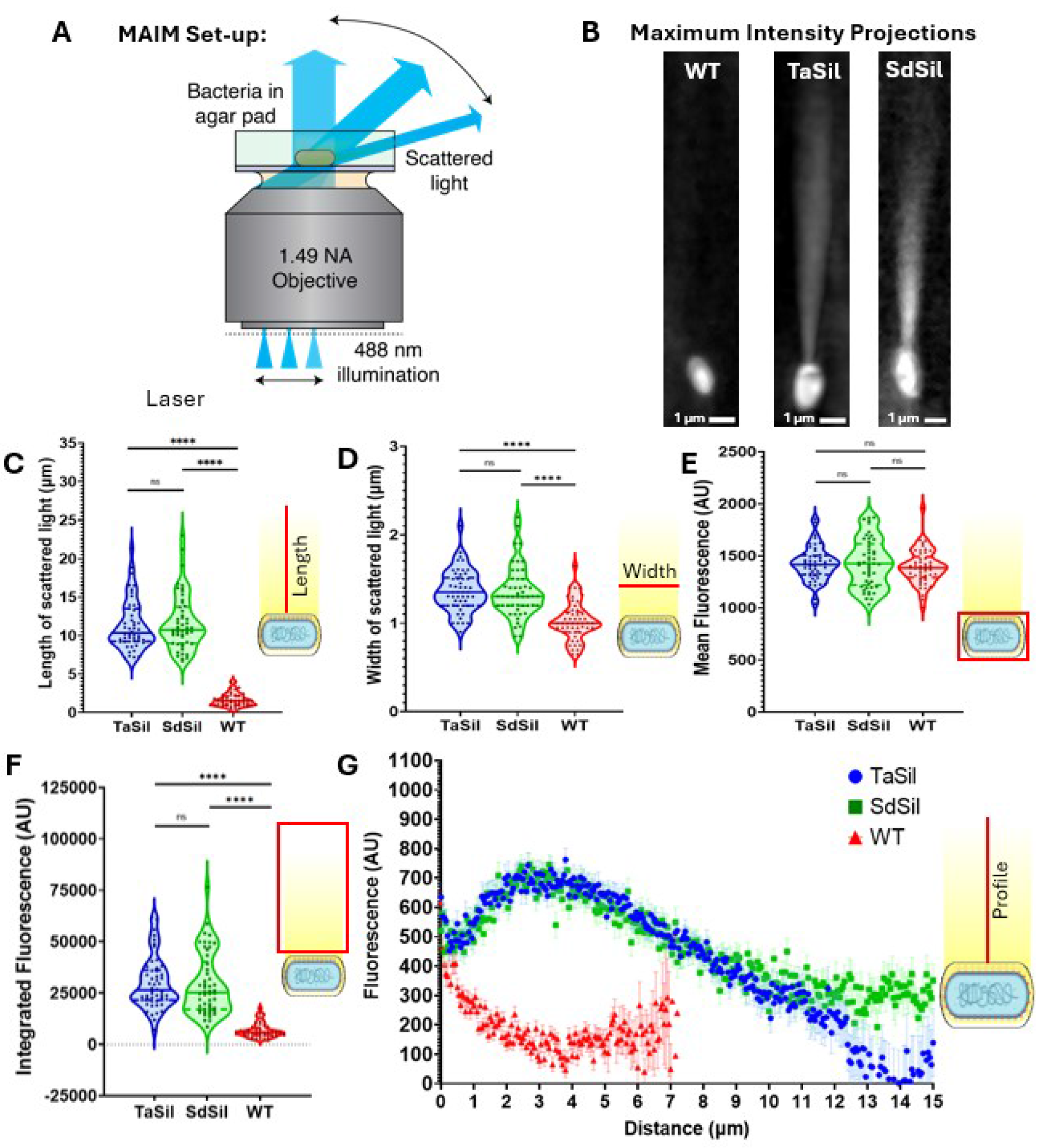
Multiple Angle Illumination Microscopy indicates that polysilicate-encapsulated cells scatter jets of light that are focused. (A) MAIM inverted microscope imaging set-up, allowing visualization of light scattered by bacteria cells at a range of incident angles. (B) Maximum intensity projections for silicatein- expressing (TaSil and SdSil) and wild-type (WT) cells scattering light via MAIM. (C) Length of scattered light, (D) width of scattered light, (E) mean intensity of light within cell boundaries, and (F) integrated intensity of the scattered light, calculated from maximum intensity projections. (G) Intensity of the scattered light as a function of distance from the edge of the cell, calculated from maximum intensity projections. Error bars correspond to standard error of the mean. (n=50) **** P≤0.0001, ns: not significant.

Videos 1-3) and created a maximum intensity projection across the sequence (Figure 4B). Wild-type cells occasionally produced diffuse, dimmer jets of scattered light, but most wild-type cells did not scatter a measurable jet of light (Figure 4B). The jets of light scattered by the encapsulated cells were significantly longer and wider than the light scattered by the wild-type cells (Figure 4C and D). The encapsulated cells and the wild-type cells showed similar illumination intensities within their cell boundaries, likely indicating some dye accumulation on the surface of both types of cells and/or autofluorescence (Figure 4E). Integration of the maximum intensity of scattered light at all illumination angles indicated that the light scattered by the encapsulated cells was significantly more intense than for the wild-type cells (Figure 4F). Analysis of the intensity of scattered light as a function of distance away from the bacterial cells revealed that the light scattered by the encapsulated cells increased in intensity with increasing distance from the cell for the first 2-3 μm, then steadily diminished in intensity thereafter (Figure 4G), indicating that the nanojets peaked outside of the encapsulated cells. By contrast, light scattered from wild-type cells showed diminishing intensity with distance from the cells with no clearly defined peak. The intensity of scattered light was more intense for the encapsulated cells than the wild-type cells at every distance from the cells (Figure 4G). These data indicate that the polysilicate-encapsulated cells are functioning analogously to dielectric microspheres, creating photonic nanojets that are focused downstream of the cell body. Encapsulated cells that had been stored for 6 months, which no longer showed metabolic activity via alamarBlue assays (Supp.Figure 11), still demonstrated scattering of nanojets that were longer, wider, and higher-intensity than the 6-month wild-type cells, with a focal peak several micrometers away from the cells (Supp. Figure 11). These results indicate that polysilicate-encapsulated cells display robust light- focusing ability, which persists for several months post-encapsulation, even after the point when the cells cease to be metabolically active.

### Encapsulated cells are metabolically active

To determine the effect of polysilicate encapsulation on bacterial physiology, we evaluated the ability of the encapsulated cells to undergo cell division as well as their metabolic activity. To investigate the growth of the silicatein-expressing cells during the induction and encapsulation process, growth curves were measured. Both the TaSil and SdSil strains exhibited slower overall growth compared to wild type, with the TaSil strain approaching zero growth while the SdSil strain showed decreasing cell density, perhaps indicating moderate toxicity of the OmpA-*S. domuncula* silicatein expression (Supp. Figure 12). To determine the effect of polysilicate encapsulation on cell division activity, the strains were induced and incubated with orthosilicate, after which the abilities of the cells to divide were measured via colony forming unit (CFU) assays. Post-encapsulation, the silicatein-expressing strains showed dramatically lower CFU/mL values compared to untreated wild-type cells, with TaSil showing approximately a five-log reduction and SdSil showing approximately a six-log reduction (Figure 5A). Upon storage, the CFUs of all strains steadily diminished over time. CFU/mL values dropped to undetectable levels after four months for the SdSil strain and after five months for the TaSil and wild-type strains.

**Fig. 5:**
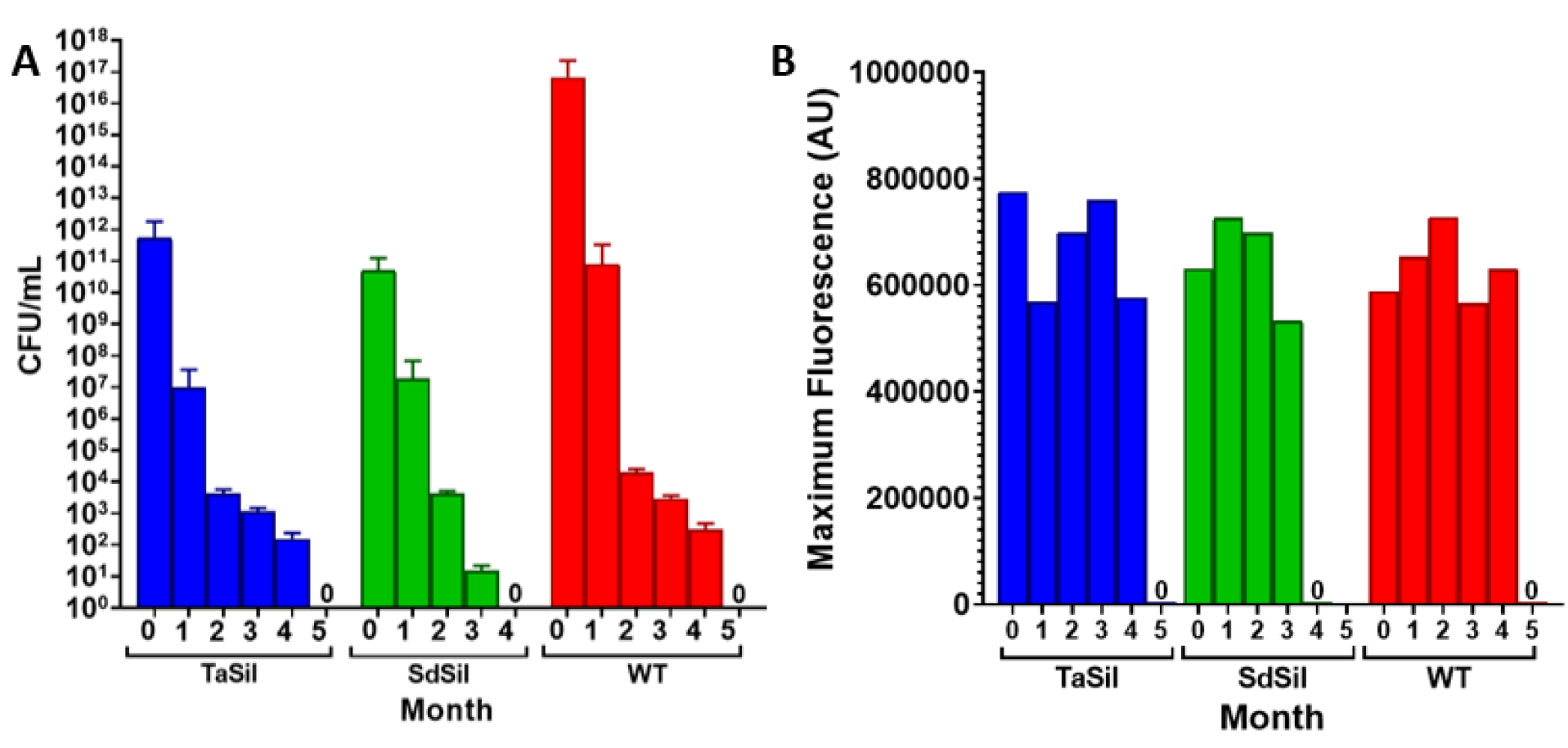
Reduced cell division and high metabolic activity of encapsulated cells. (A) CFU assay results and (B) maximum fluorescence values for alamarBlue metabolic activity assays measured for silicatein-expressing (TaSil and SdSil) cells following induction and polysilicate encapsulation and untreated wild-type (WT) cells over 5 months of storage.

Control samples of TaSil and SdSil strains that were induced but not incubated with orthosilicate showed CFU values over time that were similar to the wild-type strain, as did the empty vector control strain (Supp. Figure 13A). These experiments indicate that the polysilicate-encapsulated cells have predominantly ceased to undergo cell division.

To investigate the effect of polysilicate encapsulation on bacterial metabolic activity, the strains were analyzed using the alamarBlue assay (*31*), which produces a fluorescent signal upon interaction with the reducing cytoplasmic environment of live, metabolically-active cells.

Immediately following induction and encapsulation, TaSil and SdSil strains showed alamarBlue activity that was similar to or higher than wild-type cells (Figure 5B). Upon storage, the alamarBlue assay indicated robust intracellular reducing activity for both polysilicate- encapsulated and wild-type cells for several months, until eventually no signal was detected after four months for the SdSil strain or five months for the TaSil and wild-type strains, in agreement with the CFU assays. Silicatein-expressing strains that were induced but not incubated with orthosilicate and empty vector strains all showed similar alamarBlue activity to the wild-type strain (Supp. Figure 13B). These data indicate that polysilicate encapsulation has little effect on the metabolic activity of the bacteria cells and that the encapsulated cells remain metabolically active for several months, despite the massive reduction in cell division.

## Discussion

In this work, we demonstrate for the first time that microbes can be rationally engineered to improve their ability to focus light into photonic nanojets. We have fused the sea-sponge enzyme silicatein to the outer membrane protein OmpA, directing silicatein to the surface of *E. coli* cells where it can mineralize a polysilicate shell. In agreement with multiphysics simulations, these polysilicate-encapsulated cells are able to scatter an intense beam of light that is focused a short distance downstream from the cell, creating photonic nanojets that are much brighter than wild-type cells. Our self-assembled polysilicate-encapsulated bacteria represent the first engineered biological microlenses and serve as a proof-of-concept that cells can be engineered to act as tunable photonic components.

Our work builds on previous research using biological approaches for the production of polysilicate materials. Both sea-sponge silicatein enzymes and the silaffin peptide from diatoms have been used to create polymerized silica and silicone materials *in vitro* (*8, 32, 33*). *E. coli* bacteria have been modified to express silicatein, both cytoplasmically^7^ and on the bacterial surface, and were able to mineralize polysilicate or polylactic acid (*34*). In a separate study, *E. coli* bacteria were also modified to express recombinant silaffin R5 peptide from diatoms (*35, 6*), which was able to create silica nanostructures when co-expressed with post-translational modification enzymes that enhance the biosilification activity of the peptide. Our work is the first to demonstrate that engineered bacteria can become encapsulated in a layer of polysilicate, and that this coating enhances the optical properties of the bacteria. Furthermore, this polysilicate-mineralization activity can be implemented by introducing a single enzyme, without requiring additional, exogeneous post-translational modification enzymes.

Microparticles capable of producing photonic nanojets can be manufactured using non-biological approaches, but current techniques have several limitations. While microspheres are commercially available using materials such as silica and polystyrene, the spherical geometry is known to produce short nanojets limited to distances close to the particle surface (*37*). Such short-range nanojets make it difficult to couple the nanojets to other devices or surfaces, limiting their usefulness. Other geometries, such as microcuboids (*38*), micropyramids (*39*), and microdisks (*40*), have been explored in hopes of producing longer nanojets. Additionally, microspheres with multiple layers have been predicted to produce longer nanojets (*41*), and microspheres etched with a concentric ring pattern were shown to produce long-working-distance nanojets (*42*) . However, producing these more complicated structures has proven challenging and typically requires either low-throughput techniques (*42*) like focused ion beam etching, toxic chemicals such as hydrofluoric acid (*39*), or both.

Our approach overcomes several limitations of traditional microparticle manufacturing methods. Bacteria naturally adopt a rod-shaped geometry that is similar to a microcylinder. Although the length of these cylinders is disperse (∼1-3 µm), their width is tightly regulated and is maintained to tolerances of approximately 10% (*43*). We have shown that we can produce a layered structure by polymerizing a polysilicate shell around our bacteria, and that this layered structure extends the intensity and working distance of the associated photonic nanojets compared to uncoated bacteria. Furthermore, our microparticles do not require expensive specialized equipment to fabricate and are made under ambient conditions without the use of harsh or toxic chemicals.

Silicatein-displaying bacteria therefore offer an environmentally-friendly, high-throughput alternative to traditional manufacturing techniques.

This study provides a proof-of-concept demonstration that synthetic biological manufacturing platforms such as ours have the potential to be employed for the fabrication of a variety of different types of optical devices. Our cells have a demonstrated ability to maintain metabolic activity for an extended timeframe of four to five months post-encapsulation, opening the door to a variety of applications wherein the live bio-microlenses could be used in sense-and-response applications to respond to environmental cues by activating a reporter pathway that would change their optical properties. Additionally, the well-documented ability of the silicatein enzyme to mineralize a range of chemical substrates (*6–8, 33–35, 44–48*) could allow for the creation of bacteria coated in a variety of materials that will convey unique optical or mechanical properties. Lastly, bacteria can be engineered to grow to different sizes and shapes, offering the potential to fabricate devices with bespoke optical properties, e.g. optimizing the size and length of photonic nanojets. The application of synthetic biology to create advanced photonic devices has the potential to greatly advance fields including microscopy, nanolithography, and biomedicine (*3*).

## Materials and Methods

### Modeling of light scattering

To simulate the passage of light through the bacteria, the finite-difference time domain (FDTD) method was implemented through the use of the commercial software ANSYS Lumerical FDTD: 3D Electromagnetic Simulator. In this method, the simulation region was divided into a fine rectangular grid, and Maxwell’s equations were solved within each grid element at each discrete step in time. At one time step, the electric field was calculated, and that solution for the electric field was then used to calculate the magnetic field at the next time step. This process repeated continually, alternating between electric and magnetic field calculations until a steady state was achieved.

Within the simulations, a single bacterial cell was modeled as a rod: a cylinder whose circular faces were each adjoined to a hemisphere of equal diameter. The total length of this structure was 1.8 μm, and the diameter was 0.8 μm. For encapsulated cells, the outer polysilicate layer was modeled to be 30 nm thick by overlaying a second rod concentric with the bacterial cell, with a uniformly larger diameter of 0.86 μm. The refractive indices of the inner rod, outer rod, and surrounding background medium were set to 1.37 (*49–51*), 1.47, and 1.33, representing cytoplasm, polysilicate, and water, respectively. The cell was positioned centrally along the y- and z-axes, and its long axis was aligned at a variable angle θ to the x-axis. A plane wave of wavelength 488 nm was injected from the leftmost yz-boundary of the simulation region, propagating along the positive x-direction towards the cell. The dimensions of the simulation region were 42 μm × 6 μm × 6 μm, allowing room for the beam of scattered light to be produced. Perfectly matched layer boundary conditions were applied to all boundaries of the simulation region.

Simulations were performed separately for encapsulated cells and wild-type cells, for values of θ ranging from 0° to 90° in increments of 5°. The inclusion of a range of θ values models the random orientation of cells within the experimental methodology.

### Analysis of modeling data

A 2-dimensional intensity map was produced from the steady-state solution by summing the electric field intensity values along the range of z-values spanned by the bacterial cell. The background intensity value for each intensity map was calculated from the mean intensity of the lower-left corner square region comprising an area of 1 μm^2^, close to the light source and unaffected by the interactions between the light source and the cell. The beam region was defined as all points within the intensity map whose x-values were beyond the right edge of the cell, for which the intensity values were at least 30% greater than the background intensity value, and which were part of a contiguous set of points in the central y-region. The width of the beam was measured as the greatest difference in y-values between any two points with matching x-values along the perimeter of the beam region. The length of the beam was measured by subtracting the x-position of the right edge of the cell from the x-position of the furthest-right point within the beam region. A map of background-adjusted intensity values was produced by subtracting the background intensity value from all points within the intensity map, and any resulting negative intensity values were set to a value of 0. The intensity values were then normalized by applying the equation: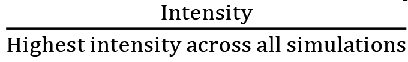 The integrated intensity of the beam was calculated by summing all intensity values within the beam region. The profile of the beam was determined by measuring the background-adjusted intensity as a function of the distance from the right edge of the cell along the centerline of the beam, defined as the subcollection of points within the beam region whose y-values were 0. To calculate the FWHM, the transverse beam profile was generated by identifying the z-position of the maximum intensity within the 3-dimensional simulation region, extracting the xy-plane of intensity values at that z-position, and measuring the intensity along the y-direction at the x-position of the maximum intensity point. The FWHM of the beam was measured as the width of the peak of the transverse beam profile at the intensity value halfway between the maximum intensity and the background intensity value of the corresponding xy-plane.

In order to approximate the effect of imaging the excitation pattern produced by the nanojet on a widefield fluorescent microscope, we convolved the results of the multiphysics model with a Gaussian approximation to the point spread function(*52*). The wavelength was estimated as 517 nm, corresponding to the emission peak of the Alexa Fluor 488 dye. The NA of the objective was set to 1.49.

### Strain information

The strain background used for all experiments is Top10 *Escherichia coli* (genotype: F- mcrA Δ(mrr-hsdRMS-mcrBC) φ80lacZΔM15 ΔlacX74 recA1 araD139 Δ(araleu) 7697 galU galK rpsL (StrR) endA1 nupG). cDNA encoding silictein genes from two species of sea sponges, *Tethya aurantia* (*21*) (GenBank: AF032117.1) and *Suberites domuncula* (*19, 21*) (GenBank: AJ272013.1), was *E. coli* codon-optimized and used to construct silicatein-expressing plasmids. OmpA-silicatein constructs were created by fusing the signaling peptide and the first nine N-terminal amino acids of lipoprotein (Lpp) from *E. coli*; amino acids 46-159 of OmpA which form its transmembrane domain (*53*); and silicatein cDNA. The plasmid containing the OmpA-*T. aurantia* silicatein (TaSil) fusion protein contains a strong ribosome binding site (B0034 (54, *55*)) upstream of the TaSil fusion protein and the rrnB T1 and T7Te terminators. OmpA-TaSil silicatein was inserted into vector pBbS5a-RFP (*56*) (Addgene #35283) replacing the vector’s RFP gene, placing TaSil expression under the control of an inducible Lac promoter (*57*). The OmpA-*S. domuncula* silicatein (SdSil) fusion protein was placed behind RBS (B0034) and cloned into the pRHA113 vector’s multiple cloning site using XbaI and BamHI, placing the SdSil gene under an inducible rhamnose promoter. The pRHA113 OmpA-*S. domuncula* silicatein construct needs to be retransformed periodically into *E. coli*, as the silicatein expression becomes more variable with longer storage at -80°C. The empty vector (EV) control in the paper refers to the background strain containing the pRHA113 vector backbone with no gene insertion, and wild type (WT) refers to the background strain containing no plasmid.

### Culture growth conditions

An overnight culture was prepared for each strain, via growth at 37°C in Luria-Bertani (LB) media on a rotator. After overnight growth, a fresh culture was inoculated via a 1/100 dilution of overnight culture into fresh LB media, with 100 µg/mL final concentration of ampicillin for TaSil, SdSil, and EV strains, in an Erlenmeyer flask. Cells were grown for 2 hours at 37°C with shaking at 250 rpm. Cells were induced by adding isopropyl ß-D-1-thiogalactopyranoside (IPTG; CAS: 367-93-1) to 1 mM final concentration for the *T. aurantia* silicatein strain (TaSil) or rhamnose to 0.2% final concentration for *S. domuncula* silicatein (SdSil) and pRHA113 empty vector (EV) strains. Cells were induced for 3 hours at 37°C with shaking, then sodium orthosilicate (Na4SiO4, Alfa Aesar, CAS: 13472-30-5) was added to a final concentration of 100 µM. Cells were incubated with silicate for 3 hours at 37°C with shaking at 250 rpm. Cells were then pelleted at 4500 rpm for 10 minutes in a swinging bucket centrifuge, and excess silicate was removed by washing three times with 1X Tris buffered saline (TBS: 50 mM Tris-HCl, 150 mM NaCl, pH 7.5). Finally, cells were resuspended in 1X TBS and stored at 4°C with constant rotation (Benchmark Scientific Roto-Mini Plus Variable Speed Rotator R2024) at 30 rpm until use.

### Cell fixation for immunofluorescence

After growth and incubation with inducer chemicals and sodium orthosilicate, cells were fixed in 2.5% glutaraldehyde in 1X phosphate buffered saline (PBS – MP Biomedicals, CAS: 2810306) for 1.5 hours at room temperature. Cells were washed three times with 1X PBS and resuspended in 1X PBS. Fixed cells were stored in 1X PBS at 4°C with constant rotation until use in immunofluorescence.

### Immunofluorescence

Fixed cells were blocked with 2% BSA in 1X PBS for 10 minutes at room temperature. Primary antibody (Silicatein alpha/silica-G antibody, Antibodies-online, ABIN191460) was added at a 1:100 dilution to the 2% BSA-cell solution. Cells were incubated with primary antibody overnight at 4°C with constant rotation. Next, cells were washed three times with 1X PBS and then incubated with the secondary antibody (Goat Anti-Rabbit IgG H&L, Alexa Fluor® 647, Abcam, ab150079) at a 1:500 dilution in 2% BSA in 1X PBS for 1 hour at room temperature in the dark. Cells were then washed three times with 1X PBS and prepped for imaging via resuspension in VectaShield® Antifade mounting medium with DAPI (Vector Laboratories UX- 93952-24). Cells were pipetted onto a 1% agarose pad and sealed with a coverslip for imaging on a Nikon A1R HD Laser Scanning Confocal Microscope using a 60X oil Apochromat TIRF objective (1.49 NA). DAPI staining of DNA was imaged using a 405 nm excitation laser and a 425-475 nm emission filter, and the secondary antibody was imaged using a 647 nm excitation laser and a 663-738 nm emission filter. Images were analyzed using Fiji software (*58*).

### Rhodamine123 staining

After growth and incubation with inducer chemicals and sodium orthosilicate, live cells were stained with Rhodamine123 (Invitrogen R302) at room temperature and in the dark where possible. 1/10 volume of 500 µM Rhodamine123 was added to the bacteria culture (i.e., 1 µL Rhodamine123 for every 10 µL culture), followed by a 15-minute incubation. Cells were washed three times with 1X TBS, then resuspended in 1X TBS. Cells were pipetted onto a 1% agarose pad and sealed with a coverslip for imaging on a Nikon A1R HD Laser Scanning Confocal Microscope using a 60X oil Apochromat TIRF objective (1.49 NA). Rhodamine123 signal was imaged using a 488 nm excitation laser and a 500-550 nm emission filter. Images were analyzed using Fiji software.

### Cell preparation for Transmission Electron Microscopy (TEM)

After growth and incubation with inducer chemicals and sodium orthosilicate, cells were fixed in a solution of 2.5% glutaraldehyde in 0.1 M sodium cacodylate buffer overnight at 4°C with constant rotation. Cells were spun down and rinsed twice for 10 minutes in the same fixation buffer. The post-fixation samples were resuspended in 1% osmium tetroxide and incubated for 30 minutes. Cells were spun down and rinsed twice for 10 minutes in dH2O, and the supernatant was removed from the pelleted cells. Cells were trapped in 3% agarose, which was cut into 1 mm cubes. The cubes were dehydrated in a graded ethanol series (50%, 65%, 80%, 95%, 100%) three times for 20 min at each step. Samples were transferred into propylene oxide:ethanol [1:1] and incubated for 30 minutes, then incubated twice for 30 minutes in 100% propylene oxide.

Samples were incubated in epoxy resin:propylene oxide [1:1] with rotation for 2.5 hours, then incubated in 100% epoxy resin overnight. Samples were embedded into molds and polymerized at 65°C for 48 hours. Cells were cut using a Leica UC7 ultramicrotome, on a diamond knife, into 1-micron sections. The sections were stained on a glass slide with Toluidine blue and rinsed with dH2O. Sections were selected for thin sectioning via light microscopy, cut to 70 nm sections, and placed onto formvar/carbon nickel slot grids (Electron Microscopy Sciences, Hatfield, PA).

Grids were stained with aqueous 2% uranyl acetate and 3% lead citrate. Cells were imaged on a Hitachi 7650 Transmission Electron Microscope at 80 kV with an attached Gatan Erlangshen 11- megapixel digital camera.

### Cell surface roughness calculations

Using Fiji, individual cells were isolated from the TEM micrographs, and the cell images were duplicated. For each cell analyzed, a 2-pixel Gaussian blur was performed on one duplicate, and a 10-pixel Gaussian blur was performed on the other. The outer edge of the cell was outlined using the lasso tool, and the perimeter was measured for both blurred images. Cell surface roughness was calculated as: Perimeter(Blur2) – Perimeter(Blur10).

<u>Scanning electron microscopy-energy dispersive X-ray spectroscopy (SEM-EDS) analysis</u> Samples stored in 1X TBS buffer were centrifuged, pelleted, and washed twice with distilled water. Each sample was resuspended in approximately 1 mL of distilled water to achieve a homogenous mixture. To prepare for SEM-EDS analysis, 50-60 µL of each sample was deposited onto an SEM sample holder and allowed to dry overnight. The samples were sputter coated with a 3 nm layer of platinum. SEM imaging was performed on a Hitachi SU3500 in secondary electron mode at an accelerating voltage of 10 keV and a working distance of 5 mm. EDS analysis was performed with an accelerating voltage of 15 keV and a working distance of 10 mm.

### X-ray powder diffraction (XRD) analysis

Samples stored in 1X TBS buffer were centrifuged, pelleted, and washed twice with distilled water. Each sample was resuspended in approximately 1 mL of distilled water to achieve a homogenous mixture. To prepare for XRD analysis, 150 µL of each sample was deposited onto an XRD sample holder with a zero-diffraction plate and allowed to dry overnight. Mineral phases of each bacterial sample were analyzed qualitatively using a Bruker D8 Advance XRD system. Cu Kα X-ray radiation with wavelength 1.5406 Å was used to scan from 10° to 60° 2θ with a step size of 0.02° and a dwell time of 1 second per step.

### Microscope setup

A custom-built microscope was used to perform MAIM. Three lasers (488 nm, 532 nm, and 640 nm) were combined into an acousto-optic modulator (Brimrose) and coupled into a single mode fiber. The other end of the fiber was coupled to a beam expander followed by a focusing lens.

These three components (fiber coupler, beam expander, and focusing lens) were all placed on a motorized translation stage (Thorlabs). The laser illumination was then reflected off a dichroic beam splitter (Chroma), allowing the beam to come to a focus at the back focal plane of a 1.49 NA oil immersion objective (Nikon), producing uniform, planar illumination at the image plane. Adjusting the motorized translation stage allowed us to continuously change the angle of illumination at the image plane. Transmitted light from an LED (Thorlabs) along with fluorescence emitted in the image plane were collected by the objective and passed through the dichroic beam splitter to a tube lens (Thorlabs). The tube lens focused the light through a second dichroic beam splitter, which separated the fluorescent emission from the transmitted LED illumination onto two sCMOS cameras (Thorlabs). This setup allowed us to image both the bacteria in brightfield and the fluorescent patterns caused by the 488 nm laser light scattered by the bacteria.

### Multiple illumination angle microscopy (MAIM)

An agarose pad was prepared for imaging by melting 1% agarose in a 0.01% poly-lysine solution and solidifying it into the desired pad shape (7/16 diameter circle, 0.5 mm thick, approximately 100 µL volume). Once solidified, the pad was placed in a dark humidity chamber and stained for 20 minutes with 20 µL of 0.5 mg/mL Alexa Fluor™ 488 NHS Ester (Invitrogen A20000) dissolved in a solution of 0.1 M sodium bicarbonate pH 8.3. Excess succinimidyl ester was quenched with a 1 M glycine rinse over the pad. Cells were then pipetted onto the fluorescently stained agarose pad and sealed with a coverslip. The cells were imaged using a custom-built inverted fluorescence microscope. Laser illumination at 488 nm was delivered to the 100X oil-immersion objective (Nikon, 1.49 NA) through a fiber launcher and telescope placed on a motorized translation stage, allowing smooth variation of the illumination angle from total internal reflection at the glass-water interface to any angle between -90° and 90° above the coverslip (where 0° corresponds to epi-illumination). Fluorescent images were acquired for multiple illumination angles spanning the full range of angles.

### MAIM image processing

MAIM images were processed using Fiji. A rolling ball background subtraction (50-pixel radius) was performed on all images and movies, then a maximum intensity Z projection was created from each movie. The maximum intensity images were used for performing image analysis including: length, width, integrated fluorescence, mean fluorescence of light within the cell boundary, and profile of scattered light. The jet beam was defined as all light scattered by a bacterial cell, for which the intensity values were greater than the background intensity value, and which were part of a contiguous area in the central region of the scattered light. The length was defined as the length of a line drawn from the edge of the cell through the center of the jet of light to the distal edge of the jet. The width was defined as the widest point of the jet beam, measured perpendicularly to the angle of incident light. The integrated fluorescence of the jet of scattered light was calculated by creating the smallest box that included the entire jet and measuring the integrated density within the area of the box. The mean fluorescence of light within the cell boundary was measured by drawing a box around only the bacterial cell and calculating the mean gray value within the box. The profile of scattered light was determined by drawing a single line through the center of the jet of light beginning at the cell edge and continuing to the first location that matched the background level of fluorescence, then graphing the profile using the plot profile function. The background level of fluorescence for each maximum intensity image was subtracted from the scattered light profiles.

### Growth curves

Cells were cultured as described above, with the exception that cultures were grown in a 96-well plate. The plate was incubated at 37°C with continuous orbital shaking at 282 rpm in a microplate reader (BioTek Synergy H1), which recorded O.D.600 measurements every 15 minutes for a total of 8.5 hours, with breaks for the addition of the inducer chemical and sodium orthosilicate.

### Colony forming unit (CFU) assay

Cells were resuspended to an O.D.600 of 0.1 in LB (with antibiotics) and were serially diluted in 1X TBS. 100 µL of the diluted cells were plated onto appropriate media for each condition (LB for WT, LB+ampicillin for plasmid-containing strains). Plates were incubated overnight at 37°C, and colonies were counted the following morning. The following equation was used to calculate the CFU/mL of the samples: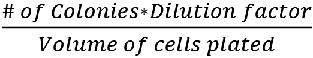

### AlamarBlue cell viability assay

After growth and incubation with inducer chemicals and sodium orthosilicate, cells were resuspended to an O.D.600 of 0.1 in LB (with antibiotics). The cells were diluted to 10^-1^ in 1X TBS. Two 100 µL aliquots of each sample were loaded into a black, clear bottom 96-well plate. Half of the samples were monitored for alamarBlue fluorescence (Ex: 530 nm/Em: 590 nm) and the other half for absorbance at 600 nm to determine a growth curve, since the absorbance spectrum of alamarBlue overlaps with the O.D.600 measurements used to determine growth curves. 10 µL of alamarBlue (Invitrogen DAL1025) was added to the samples that were monitored for fluorescence. Each sample and dilution were analyzed in triplicate. A BioTek Synergy H1 microplate reader was used to incubate and continuously orbital shake (282 rpm) the 96-well plate at 37°C. Readings were taken every 15 minutes for 24 hours. 1X TBS samples were used as negative controls for the alamarBlue assays, and LB samples were used as negative controls for the cell growth curves. Fluorescence or O.D.600 values of negative control samples were subtracted from each experimental data set to correct for background signal.

### Statistical analyses

Wilcoxon t-tests were performed using GraphPad Prism version 9.1.0 for Windows, GraphPad Software, San Diego, California USA for all statistical analyses.

## Supporting information

Supplemental Video 1: TaSil

Supplemental Video 2: SdSil

## Acknowledgments

The authors wish to thank TU Delft’s 2016 iGEM team for the initial development and work done on this project: Carmen Berends, Célina Reuvers, Charlotte Koster, Giannis Papazoglou, Iris de Vries, Lara van der Woude, Liza de Wilde, and Tessa Vergroesen; Filipe Natalio for initial discussions; Matthew Hamilton Fyfe from University of Colorado Boulder for assistance with elemental analyses laboratory support in Wil V. Srubar III’s laboratory; Morgan Brady from the University of Rochester for laboratory and project development and suggestions; Karen Bently, Chad Galloway, and Kelsea Cristillo of the Electron Microscopy Resource in the Center for Advanced Research Technologies (CART) at the University of Rochester Medical Center for performing electron microscopy imaging; the University of Rochester Medical Center’s Center for Advanced Light Microscopy and Nanoscopy (CALMN), especially Julie Zhang and Kaye Thomas for their assistance with confocal microscopy; Brian McIntyre and Sean O’Neil from the University of Rochester’s Integrated Nanosystems Center for electron microscopy imaging and technique guidance.

Funding to A.S.M., L.M.S., M.M.B., and E.A.A. was provided by the National Science Foundation via MODULUS DSM-2031180, ITE-2137561, and ITE-2230641, and by the National Institutes of Health via 1R01GM143182-01. L.M.S. was supported in part by a fellowship award under contract FA9550-21-F-0003 through the National Defense Science and Engineering Graduate (NDSEG) Fellowship Program, sponsored by the Air Force Research Laboratory (AFRL), the Office of Naval Research (ONR) and the Army Research Office (ARO).

I.R. and P.S.C. were funded by the University of Rochester. Funding for B.C.A., J.R., and W.V.S.III was provided by the National Science Foundation via CMMI-1943554.

## Data and material availability

All data are available in the manuscript or the supplementary materials.

## Author contributions

L.M.S. and A.S.M. conceived and planned the study. L.K., M.V.V., and L.M.S. created the DNA constructs. L.M.S. performed the microbiology characterization and microscopy analyses.

E.A.A. developed the microscopy imaging techniques. M.M.B., I.R., P.S.C., and G.R.S. performed the modeling. B.C.A., J.R., and W.V.S. performed the SEM-EDS and XRD analyses. All authors contributed to the writing of the final manuscript and authorized its publication.

