## Supplemental Information for "Engineered bacteria that self-assemble “bioglass” polysilicate coatings display enhanced light focusing"

**This PDF file includes:**

Figs. S1 to S12

**Other supporting materials for this manuscript include the following:**

Movies S1 to S3

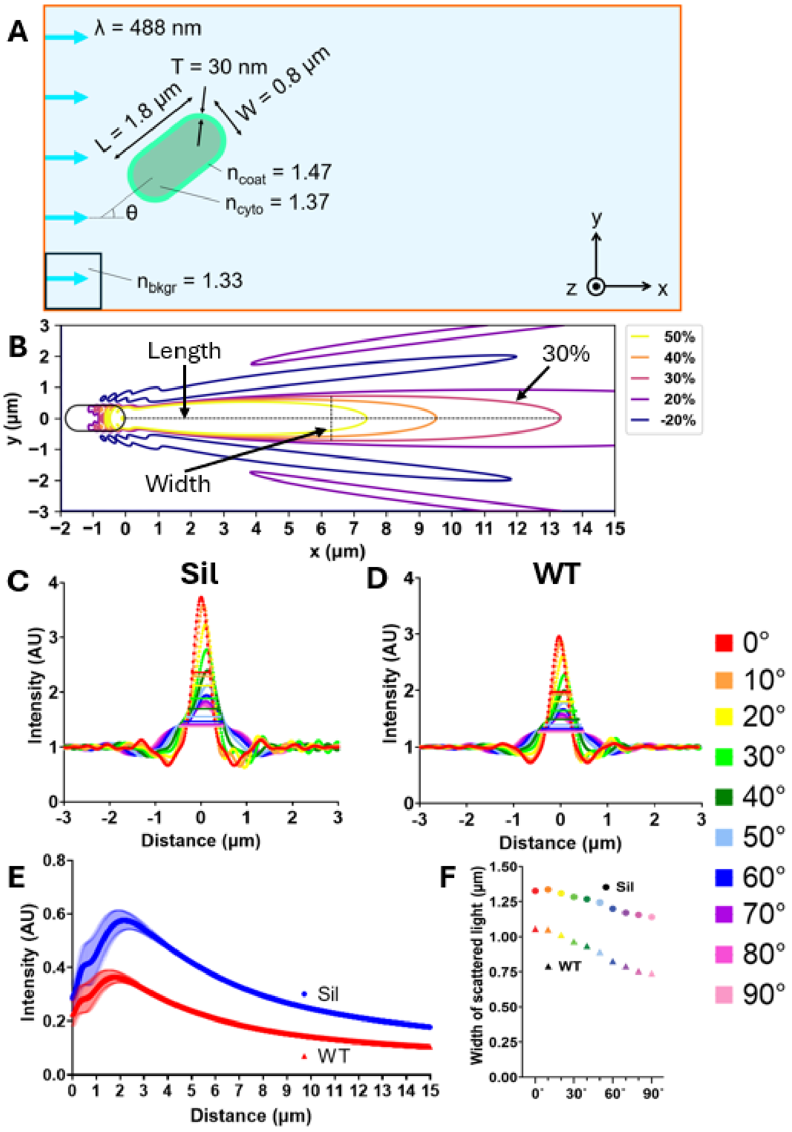


**Fig. S1: Mathematical modeling predicts that the angle of incoming light affects the intensity and focal distance of light scattered by polysilicate-encapsulated cells**

(A) Modeling parameters, not drawn to scale, used in simulations of light scattering, including the dimensions of the modeled cell and its polysilicate coating, the refractive indices of the polysilicate, the bacterial cytoplasm, and the surrounding water, and the angle and wavelength of incoming light. The black box in the lower left corner corresponds to the 1 $\mu m^2$ square used for background calculations. (B) Example contour plot of electric field intensity from a polysilicate-encapsulated bacteria cell, showing parameters used to analyze data: beam percentage above background, length, width, and cell boundary. (C-D) Transverse beam profiles, where each color corresponds to a different angle of the cell in relation to the incident light, with FWHM indicated by a horizontal line, for encapsulated cells (Sil) (C) and wild-type cells (WT) (D). (E) Intensity of the scattered light as a function of distance from the edge of the cell, averaged over results from incident light angles 0°-90° at 5° increments. Error bars correspond to standard error of the mean. (F) Width of scattered light, where each color corresponds to a different angle of the cell in relation to the incident light.

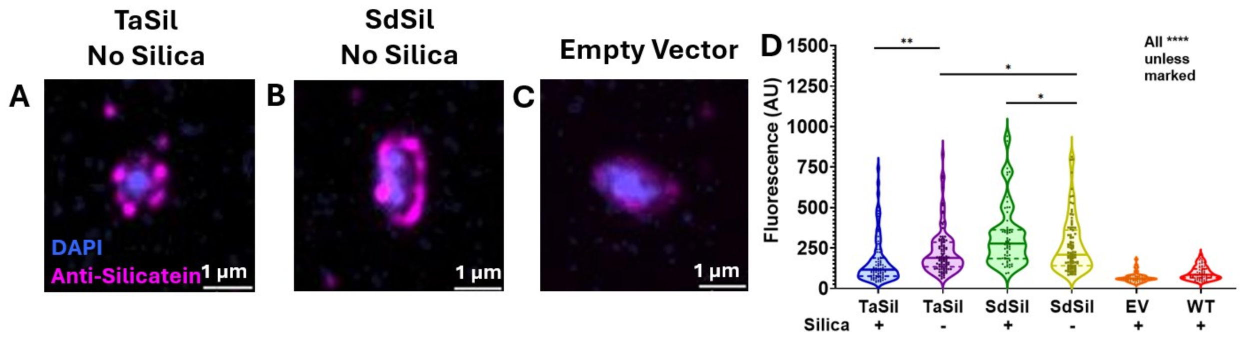


**Fig. S2: Immunofluorescence shows silicatein localization to the cell borders of silicatein-expressing cells in the absence of orthosilicate**

(A-C) Immunofluorescence of induced silicatein-expressing strains that were not incubated with orthosilicate (TaSil no silica and SdSil no silica) (A and B) and mock-induced empty vector strain incubated with orthosilicate (C). (D) Quantification of antibody fluorescence intensity of A-C, Figure 1A-C ($n_{TaSil}$=90, $n_{TaSil\ no\ silica}$=100, $n_{SdSil}$=60, $n_{SdSil\ no\ silica}$=90, $n_{EV}$=60, $n_{WT}$=60). ****$P \leq 0.0001$, ** $P \leq 0.01$, * $P \leq 0.05$.

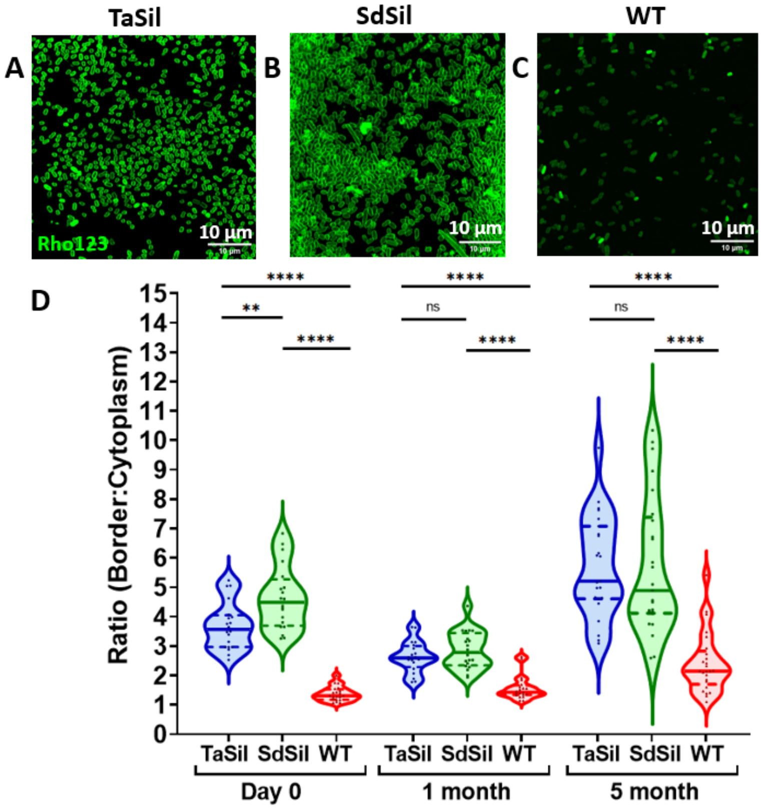


**Fig. S3: Polysilicate staining reveals a high border-to-cytoplasm ratio for silicatein-expressing cells for 5 months**

(A-C) Larger field of view of Rhodamine123-stained silicatein-expressing (TaSil and SdSil) (A and B) and wild-type (WT) (C) strains. (D) Quantification of border to cytoplasm Rhodamine123 staining for day 0 (n=20), 1-month (n=20), and 5-month samples (n=25). **** P≤0.0001, ** P≤0.01, ns: not significant.

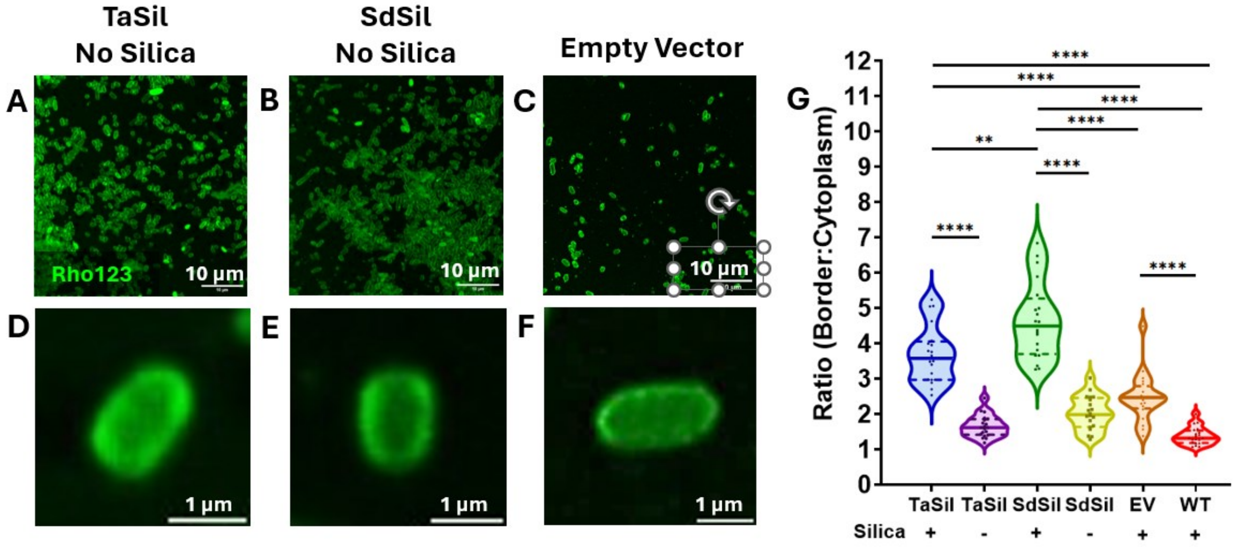


**Fig. S4: Polysilicate encapsulation requires orthosilicate and silicatein-expressing plasmids** (A-C) Rhodamine123 staining of induced silicatein-expressing strains that were not incubated with orthosilicate (A and B) and mock-induced empty vector strain incubated with orthosilicate (C). (D-F) Individual cells stained with Rhodamine123 from the samples in A-C. (G) Quantification of border to cytoplasm Rhodamine123 staining of D-F and Figure 2E-G (n=20). ****P≤0.0001, ** P≤0.01, * P≤0.05.

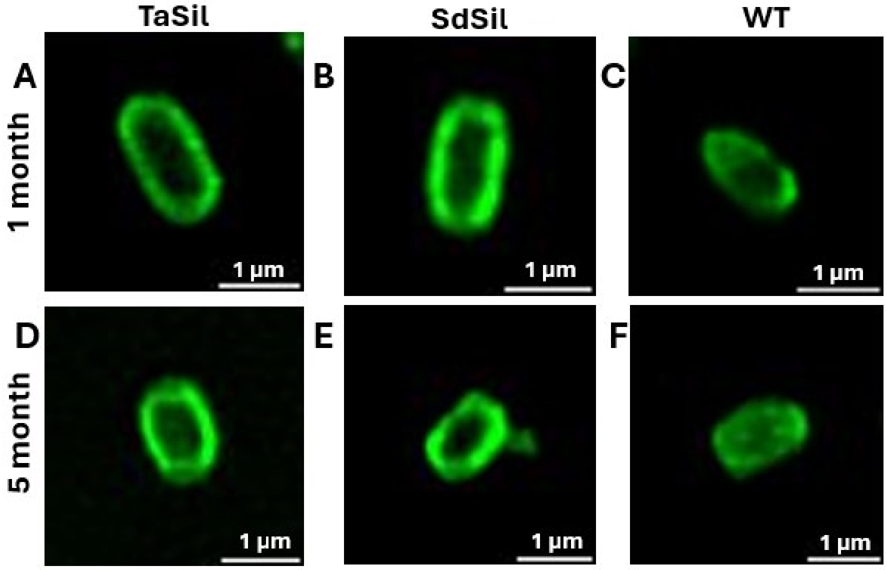


65 **Fig. S5: Rhodamine123 staining remains consistent over time**
(A-C) Rhodamine123 staining of 1-month-old samples of silicatein-expressing (A and B) and wild-type (C) strains. (D-F) Rhodamine123 staining of 5-month-old samples of silicatein-expressing (D and E) and wild-type (F) strains.

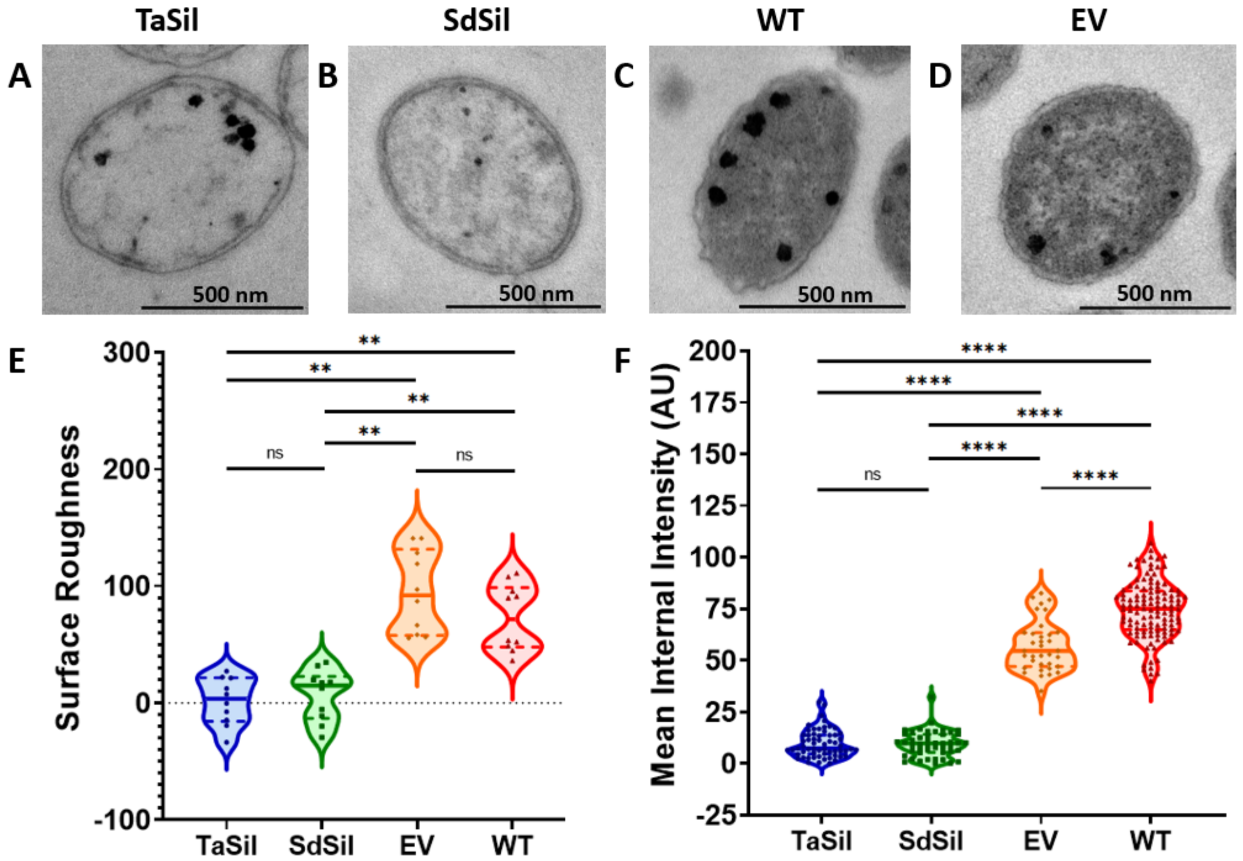


**Fig. S6: Silicatein-expressing cells have lower surface roughness**

(A-D) TEM thin section imaging of silicatein-expressing (TaSil and SdSil) (A and B), wild-type (WT) (C), and empty vector (EV) (D) strains. (E) Quantification of surface roughness of A-D (n=10). (F) Quantification of internal electron density of each strain, normalized to background electron density ($n_{TaSil}$=61, $n_{SdSil}$=47, $n_{EV}$=35, $n_{WT}$=118). **** $P \leq 0.0001$, ** $P \leq 0.01$, ns: not significant.

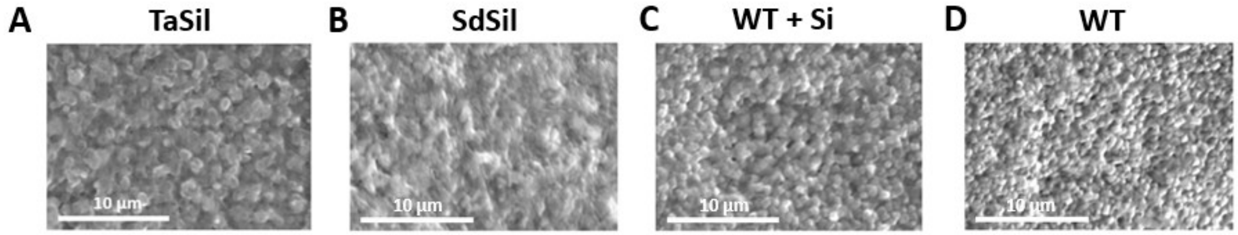


**Fig. S7: SEM images of silicatein-expressing cells used for SEM-EDS**
(A-D) SEM images of silicatein-expressing strains (TaSil and SdSil) (A and B) following silicatein induction and incubation with orthosilicate, and wild-type strains (C and D), where C was incubated with orthosilicate (WT + Si) and D was not (WT).

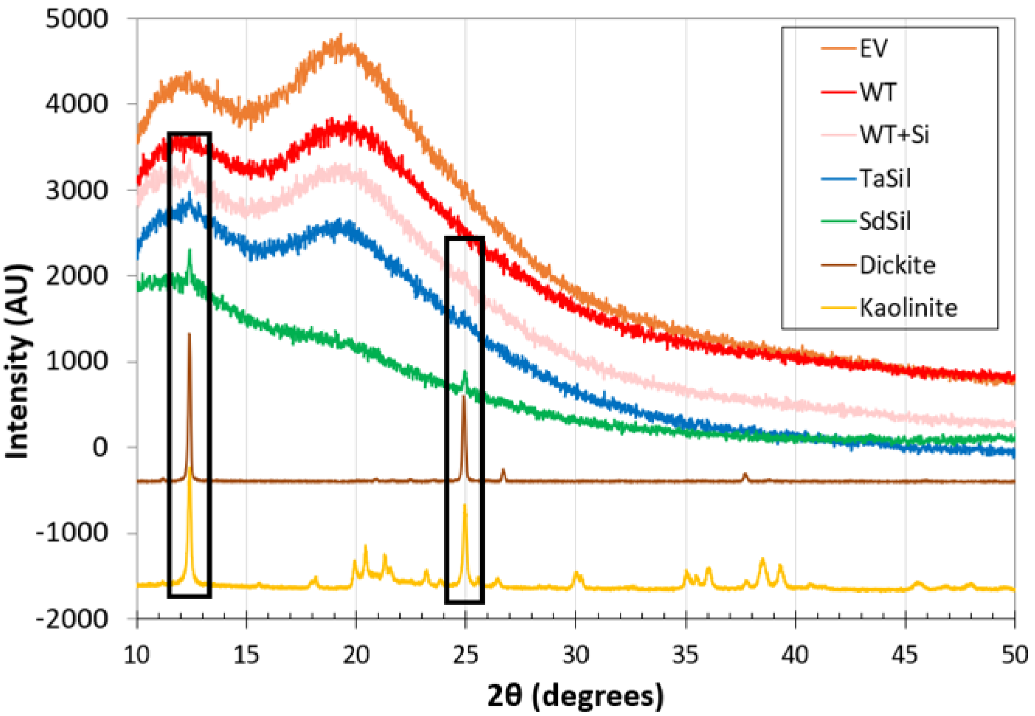


**Fig. S8: Silica detection for silicatein-expressing cells via XRD analysis.**
X-ray diffraction analysis of silicatein-expressing, polysilicate-encapsulated strains (TaSil and SdSil), wild-type strain (WT), wild-type strain incubated with orthosilicate (WT+Si), and empty vector strain (EV). Boxes correspond to peaks for kaolinite and/or dickite, which are minerals containing Al and Si.

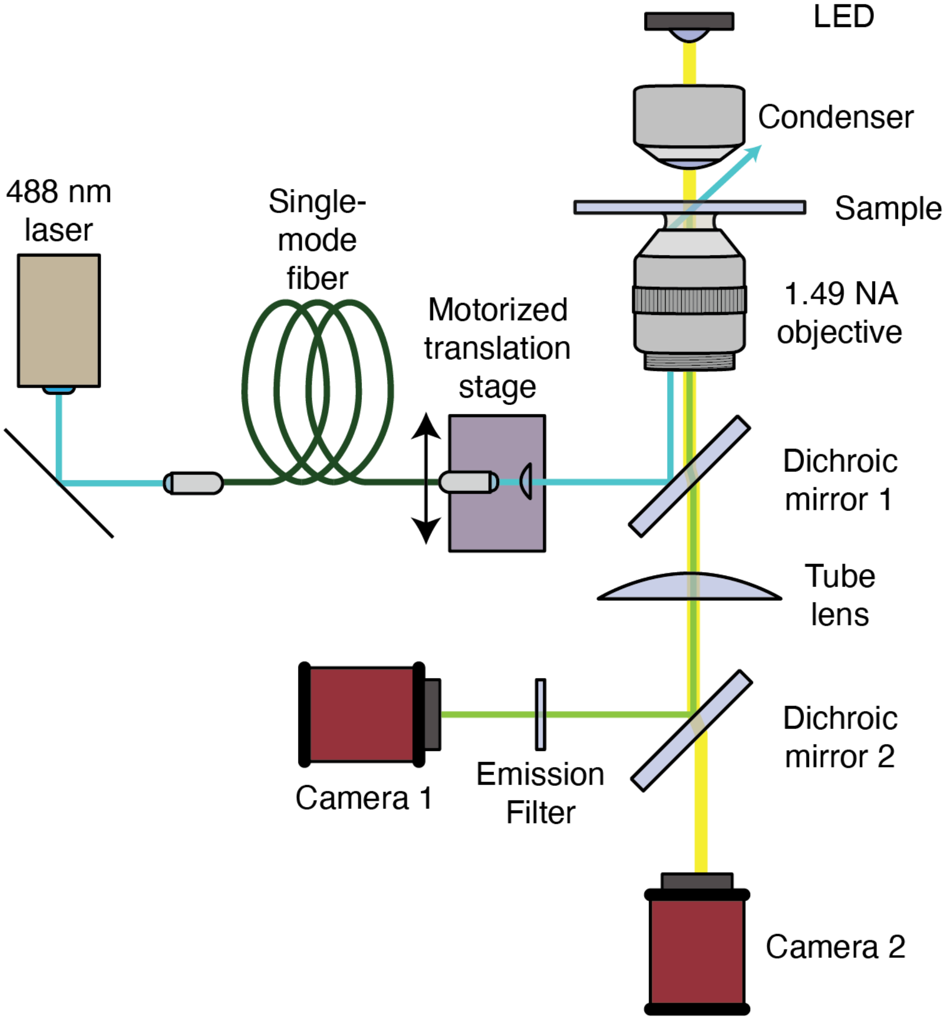


**Fig. S9: Simplified diagram of microscope used for MAIM**
MAIM illumination (light blue line) is supplied by a 488 nm laser. The beam is coupled to a single mode fiber with the opposite end placed on a motorized translation stage. The beam is expanded and focused by optics that share the same translation stage, allowing the beam to be translated laterally. The beam is then reflected by dichroic mirror 1 so that the focal point lies in the back focal plane of the high NA objective. This setup produces broad illumination in the image plane at a variable angle of incidence. Light scattered by the MAIM illumination excites fluorescent dye around the bacteria, and the emitted fluorescence (green line) is collected by the objective, passed through dichroic mirror 1, focused by the tube lens, and then directed by dichroic mirror 2 onto camera 1. A separate brightfield illumination path (yellow line) comes from a 430 nm LED focused by a condenser lens onto the sample. The brightfield illumination is transmitted through the sample and collected by the objective. From here, the light is passed through dichroic mirror 1, focused by the tube lens, passed through dichroic mirror 2, and then imaged onto camera 2.

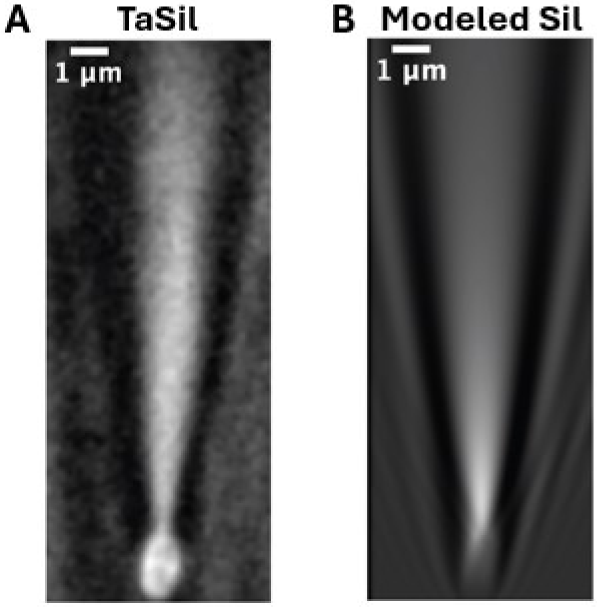


**Fig. S10: Comparison of imaged and predicted nanojets.**
(A) At nearly transverse illumination (between 85˚ and 90˚), the photonic nanojet emerging from a silicatein-expressing (TaSil) bacterium was visualized using Alexa 488 dye in solution, displaying a V-shaped pattern with two side lobes. (B) The results of the multiphysics model (for a cell angle of 10˚ to the incident light) were convolved with a Gaussian approximation to the point spread function to approximate what the microscope should detect. A similar V-shaped pattern with side lobes is again visible.

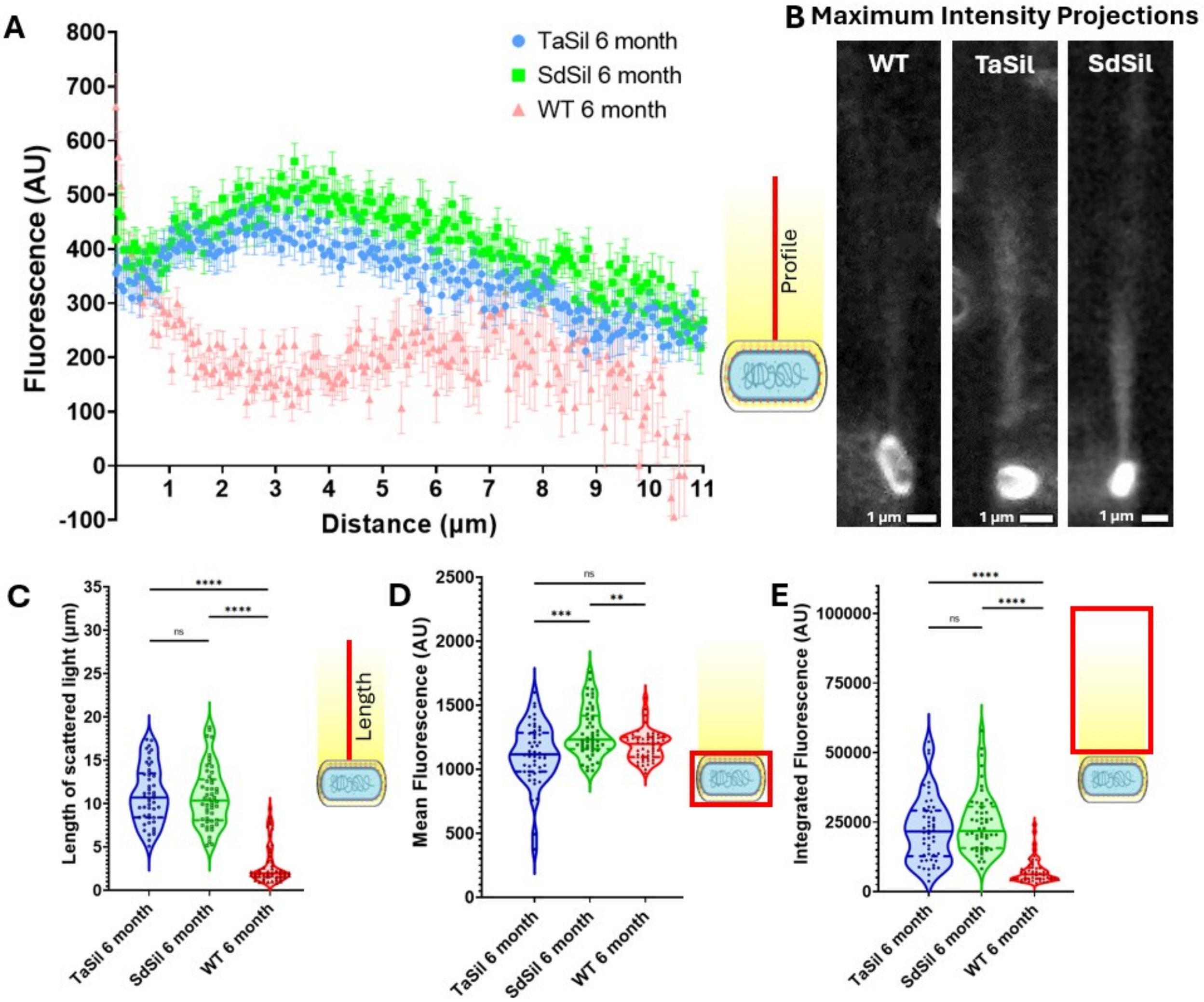


**Fig. S11: MAIM lensing of 6-month-old polysilicate-encapsulated cells**

(A) Intensity of the scattered light as a function of distance from the edge of the cell for 6-month-old silicatein-expressing (TaSil and SdSil) and wild-type (WT) cells scattering light via MAIM, calculated from maximum intensity projections. (B) Maximum intensity projections for 6-month-old silicatein-expressing (TaSil and SdSil) and wild-type (WT) cells scattering light via MAIM. (C) Length of scattered light, (D) mean intensity of light within cell boundaries, and (E) integrated intensity of the scattered light, calculated from maximum intensity projections. Error bars correspond to standard error of the mean. (n=50) **** $P \leq 0.0001$, ns: not significant.

125

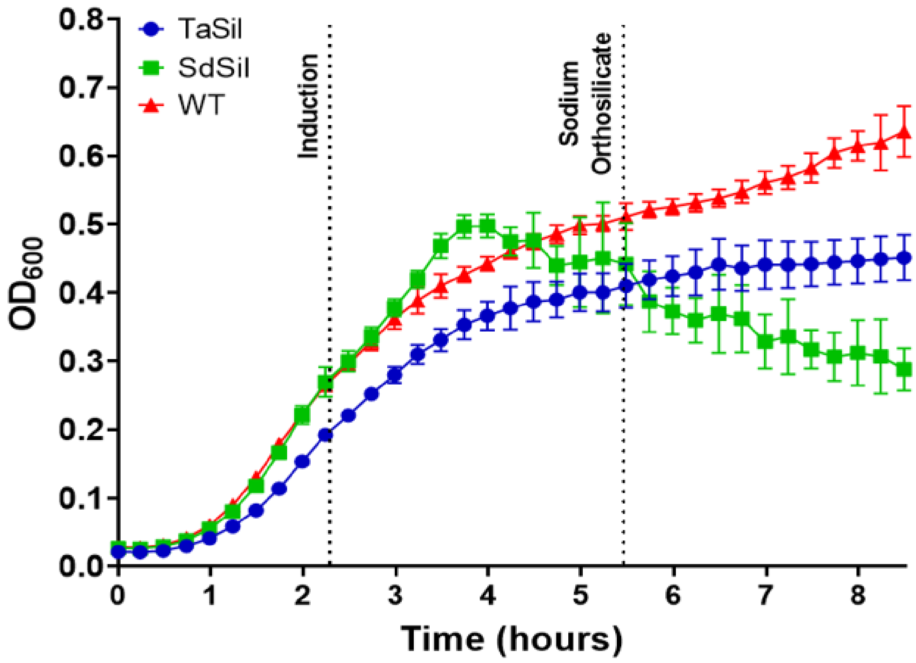


**Fig. S12: Silicatein expression slows cell division rate**
Growth curve of the silicatein-expressing (TaSil and SdSil) and wild-type (WT) strains over the time frame of the induction and encapsulation protocol. Error bars correspond to standard deviation.

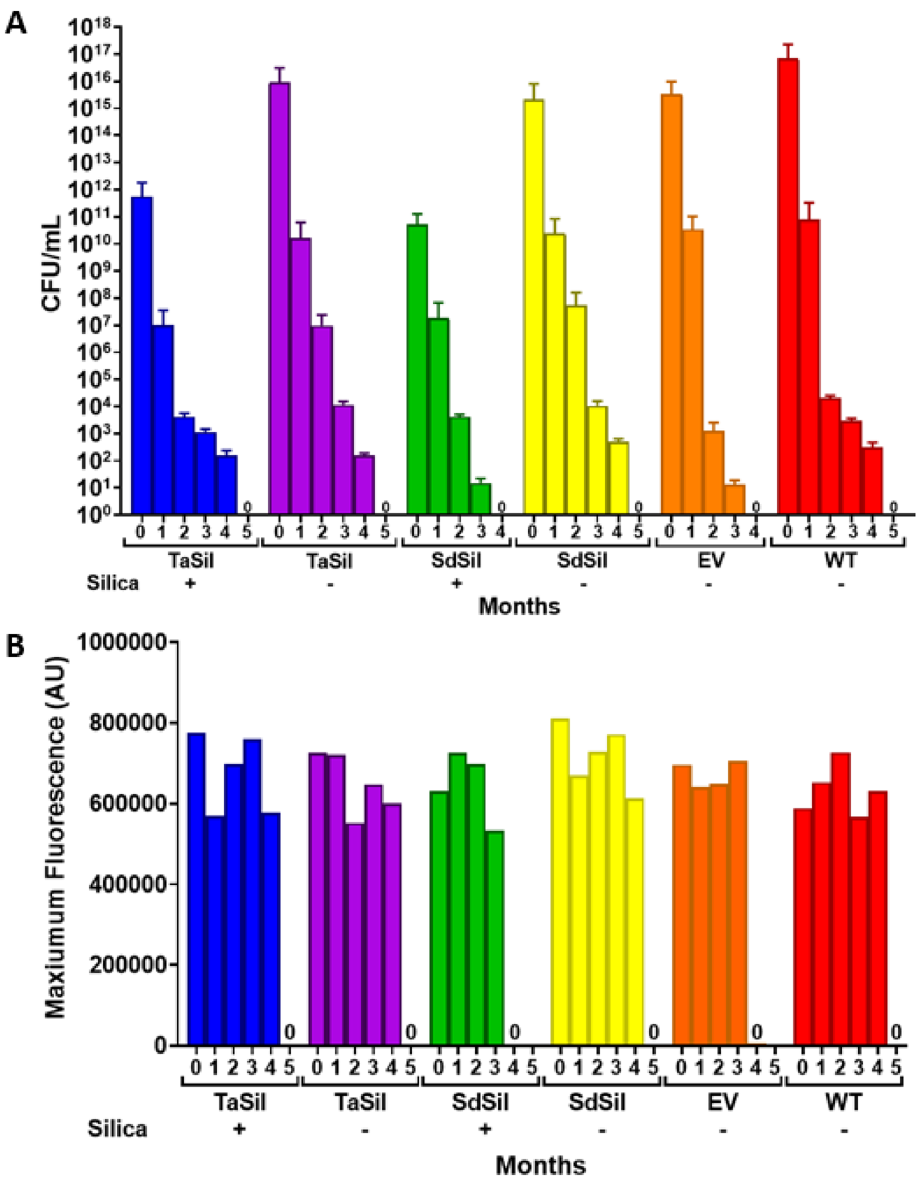


**Fig. S13: Reduced cell division of polysilicate-encapsulated cells requires orthosilicate**
(A) CFU assay results and (B) maximum fluorescence values for alamarBlue metabolic activity assays measured for silicatein-expressing (TaSil and SdSil) cells following induction, both with and without polysilicate encapsulation, as well as untreated wild-type (WT) and mock induced empty vector (EV) cells, over 5 months of storage.

**Movie S1: TaSil MAIM**
Video showing MAIM of TaSil polysilicate-encapsulated strain.

**Movie S2: SdSil MAIM**
Video showing MAIM of SdSil polysilicate-encapsulated strain.

**Movie S3: WT MAIM**
Video showing MAIM of WT strain.
